# Perturb-seq reveals distinct responses to pluripotency regulator dosages underlying the control of self-renewal and differentiation

**DOI:** 10.1101/2025.08.07.669196

**Authors:** Jielin Yan, Hyein S. Cho, Renhe Luo, Michael A. Beer, Wei Li, Danwei Huangfu

## Abstract

Precise regulation of transcription factor (TF) expression is critical for maintaining cell identity, but studies on how graded expression levels affect cellular phenotypes are limited. To address this gap, we employed human embryonic stem cells (hESCs) as a dynamic model to study gene dosage effects and systematically titrated key TFs *NANOG* and *OCT4* expression using CRISPR interference (CRISPRi). We then profiled transcriptomic changes in hESCs under self-renewal and differentiation conditions using single-cell RNA-seq (scRNA-seq). Quantitative modeling of these Perturb-seq datasets uncovers distinct response patterns for different types of genes, including a striking non-monotonic response of lineage-specific genes during differentiation, indicating that mild perturbations of hESC TFs promote differentiation while strong perturbations compromise it. These discoveries suggest that fine-tuning the dosage of stem cell TFs can enhance differentiation efficiency and underscore the importance of characterizing TF function across a gradient of expression levels.

## Introduction

The expression levels of cell type-specific transcription factors (TFs) are tightly regulated to ensure the precise control of cell identity. In development, altered expression levels of a single TF can affect the spatiotemporal regulation of lineage specification and proper activation of downstream gene programs. Many TFs exhibit haploinsufficiency in humans^1^. For example, a mutation in a single *GATA6* allele can severely impair pancreatic development, resulting in neonatal diabetes^2–4^. In complex traits and diseases, numerous studies on genome-wide associations and expression quantitative trait loci (eQTL) support that genomic variations often modulate gene expression and confer phenotypic differences in a cumulative manner^5–12^. Therefore, there is an urgent need to better understand how varying TF dosage quantitatively influences downstream gene expression and cell states.

Gene functions are primarily studied through genetic knockout or knockdown experiments. These approaches, while essential, cannot capture the full spectrum of phenotypic consequences resulting from different gene expression levels, especially those involving small variations. Phenotypic characterization across multiple dosages can be technically challenging and prohibitively labor- and cost-intensive. Recent advances have begun to ease these constraints, enabling both precise dosage manipulation and comprehensive transcriptomic profiling. Notable examples include the use of degron tags to deplete protein expression^13^ and the potential application of single-cell RNA-seq (scRNA-seq) to simultaneously profile transcriptional changes across many dosage conditions^14^. Despite these advances, systematic studies of gene dosage effects remain limited, especially in dynamic systems. We have previously shown that cells undergoing developmental cell state transitions are particularly sensitive to TF dosage^15^, highlighting the need for a quantitative understanding of how TF dosage affects downstream gene expression and differentiation outcomes. However, a major challenge of understanding gene dosage effects during differentiation is the cellular and transcriptional heterogeneity inherent to dynamic processes. To overcome this issue, we utilized Perturb-seq, which combines CRISPR interference (CRISPRi)-mediated modulation of gene dosage with scRNA-seq. This enables us to account for cellular heterogeneity while simultaneously collecting transcriptomic data from an array of dosage conditions for computational modeling.

We chose human embryonic stem cells (hESCs) as a sensitive and dynamic model to study the dosage effect of TFs on self-renewal and differentiation. Pluripotency TFs NANOG and OCT4 (*POU5F1*) allow hESCs to self-renew and to maintain differentiation potency, but their down-regulation is needed for the cells to differentiate, raising the question of how hESCs fine-tune their response to varying levels of these TFs.^16–22^. To address this, we generated different dosages of *NANOG* and *OCT4* and measured their effect by performing Perturb-seq in the context of ESC self-renewal and definitive endoderm (DE) differentiation, one of the first key cell fate transitions from pluripotency. Quantitative dosage modeling reveals distinct response patterns for different classes of genes. In particular, we found that genes associated with pluripotency and differentiation display strong switch-like responses to *NANOG* and *OCT4* dosages at the ESC stage, indicating that cell identity is regulated with high cooperativity. In DE differentiation, in addition to switch-like responses, we also observed a distinct non-monotonic response pattern, with many DE genes being first up-regulated and then down-regulated with increasing levels of perturbation. This result supports the hypothesis that mild reduction of pluripotency TF dosages promotes differentiation while strong reduction compromises it, suggesting that differentiation efficiency can be maximized by partially reducing the dosage of stem cell TFs. These discoveries not only enhance our understanding of pluripotency but also highlight the non-linearity of gene expression response to TF dosages, emphasizing the importance of characterizing TF function across a gradient of expression levels.

## Results

### Perturbations of pluripotency TFs uncover dosage-dependent rewiring of the hESC transcriptome

To achieve varying dosages of *NANOG* and *OCT4*, we designed a gRNA library based on results from a recent CRISPRi screen we conducted under hESC self-renewal conditions^23^ (Figure S1A). This screen used an array of gRNAs to target putative enhancer regions of *NANOG* and *OCT4*, among other TFs. We discovered two *NANOG* enhancers, which we designated *NANOG* e1 and e2, and also recovered the well-characterized *OCT4* proximal and distal enhancers^24–27^. We reasoned that perturbing these enhancers enabled modulation of their target genes’ expression levels. Given the well-established requirement of *NANOG* and *OCT4* for hESC self-renewal, we used depletion effects from the self-renewal screen to guide the selection of a panel of gRNAs with varying perturbation strengths (Figure S1B). We also included gRNAs targeting the promoters of *NANOG* and *OCT4* as well as negative control safe-targeting (ST) or non-targeting (NT) gRNAs (Figure 1A, Table S1). In total, we cloned 48 gRNAs into the CROP-seq backbone^28^ to make the lentiviral library for CRISPRi and scRNA-seq capture. hESCs carrying doxycycline-inducible dCas9-KRAB were infected with the library, treated with doxycycline for 6 days, and harvested for scRNA-seq (referred to hereafter as the ESC Perturb-seq dataset) (Figure 1A).

**Figure 1.**
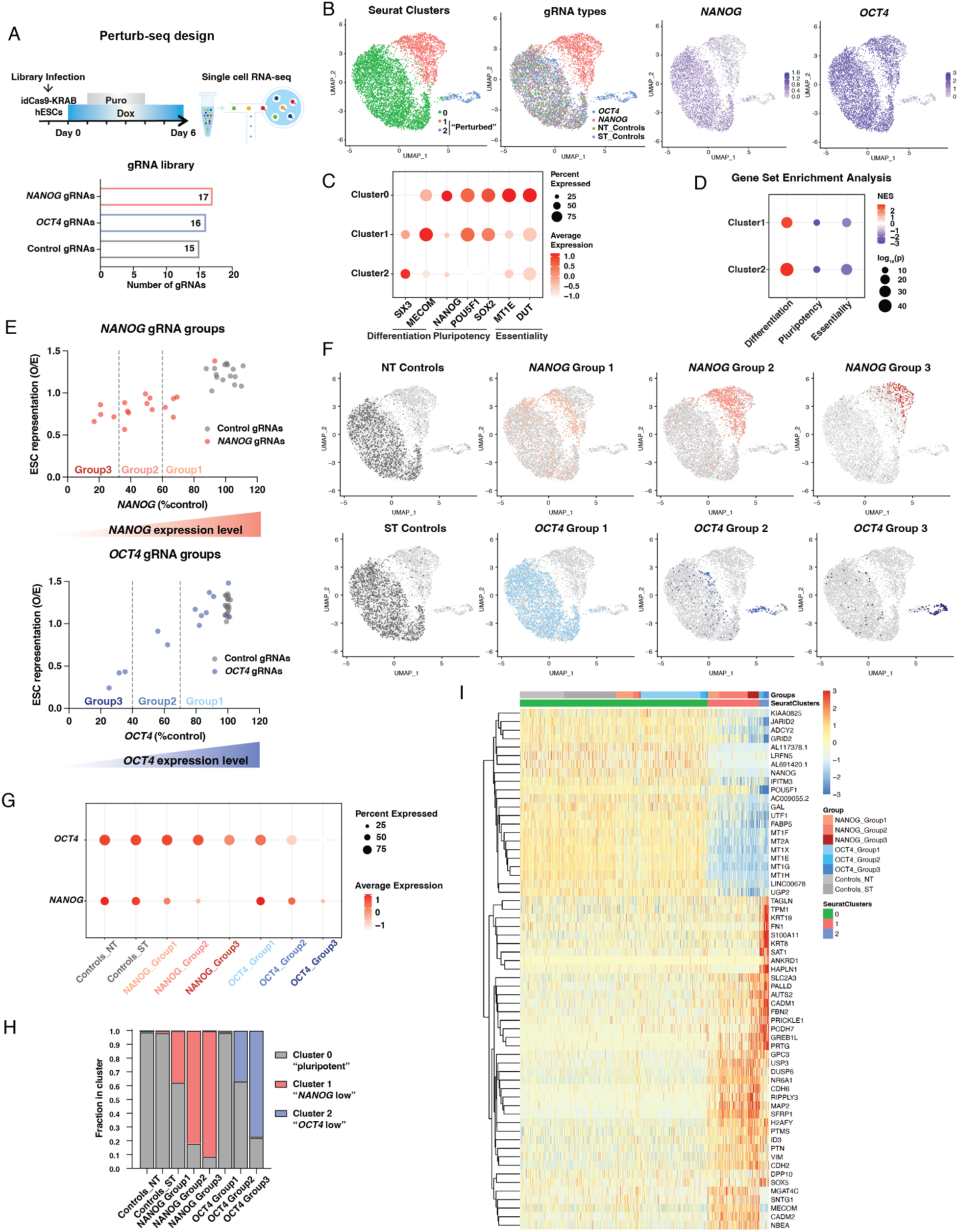
ESC Perturb-seq uncovers dosage-dependent transcriptomic rewiring to titrated perturbations. A. Schematic of ESC Perturb-seq experimental design. B. UMAPs showing Seurat clustering, gRNA distribution, *NANOG* expression, and *OCT4* expression. C. Cluster marker genes associated with differentiation, pluripotency, and essentiality. D. Summary of GSEA showing enrichment of differentiation, pluripotency, and essentiality gene sets, ranked by expression log2FC in Cluster 1 or Cluster 2 vs Cluster 0. E. Average *NANOG* and *OCT4* expression and representation for each gRNA, and expression-based groupings. F. UMAPs showing distribution of gRNAs from different perturbation groups in Seurat clusters. G. Expression of *NANOG* and *OCT4* in each perturbation group. H. Quantification of gRNA distribution in F. I. Heatmap with hierarchical clustering of top 20 marker genes from each cluster, ordered by Seurat clustering and perturbation groups. Color key indicates scaled expression.

Seurat^29^ analysis identified three clusters, with Cluster 1 enriched for *NANOG* gRNAs, Cluster 2 for *OCT4* gRNAs, and Cluster 0 showing a mixture for all gRNA types (Figures 1B and S1C). Consistent with the gRNA distribution, *NANOG* and *OCT4* expression levels were significantly reduced in Cluster 1 and 2, respectively, compared to Cluster 0 (Figures 1B-C and S1D). *NANOG*, a target gene of OCT4^16^, was also down-regulated in Cluster 2, suggesting these cells experienced both primary and secondary perturbation effects. Therefore, Cluster 1 and 2 are referred to as “perturbed clusters” hereafter, with Cluster 0 representing the normal pluripotency state. Gene ontology (GO) analysis of DEGs in the perturbed clusters revealed up-regulation of morphogenesis-related genes and down-regulation of housekeeping genes (Table S2, Figures S1E), in line with NANOG and OCT4’s roles in preventing pre-mature differentiation^22,30^, and the requirement of essentiality genes for hESC self-renewal^31^. Gene set enrichment analysis (GSEA)^32^ of the perturbed clusters further identified up-regulation of differentiation genes, defined as genes induced during DE or neuroectoderm (NE) differentiation (Figure S1F), and down-regulation of pluripotency and essentiality genes (Table S3, Figures 1C-D, S1G). Together, these results indicate cells in the perturbed clusters have shifted away from the normal ESC state.

We next examined quantitative changes of gene expression in response to varying levels of perturbation. Aggregating *NANOG* and *OCT4* expression by gRNA genotypes to determine the guides’ on-target knockdown (KD) efficiencies, we confirmed that our Perturb-seq dataset captured a range of perturbation strengths suitable for gene dosage analysis (Figure 1E). gRNAs with stronger perturbation strengths tended to be more depleted, supporting our gRNA selection based on depletion in the CRISPRi screen^23^. To stratify the perturbation effects, we grouped *NANOG* and *OCT4* gRNAs based on their on-target KD efficiencies, with Group 1, 2 and 3 corresponding to mild, intermediate and strong perturbations, respectively (Figures 1E-G, S1H). Consistent with these groupings, the number of differentially expressed genes (DEGs) increased with the perturbation strength (Figure S1I). This dosage-dependent transcriptional rewiring was also evident in the UMAP, where cells increasingly localized to Cluster 1 or 2 as *NANOG* or *OCT4* expression was progressively repressed (Figure 1H). Visualization of the top 20 marker genes from each cluster on a heatmap showed a greater degree of expression changes in cells receiving stronger perturbations (Figure 1I). Taken together, these Perturb-seq results demonstrate clear quantitative response of the hESC transcriptome to *NANOG* and *OCT4* dosages.

### Quantitative modeling reveals distinct modes of dosage response

We aimed to quantitatively characterize how cell states respond to perturbations. Specifically, we asked whether hESCs exit the pluripotent state abruptly upon reaching a threshold level of *NANOG* or *OCT4* depletion, or whether they transition gradually through a continuum of intermediate states as perturbation strength increases. To quantify cell state changes, we first calculated “perturbed” gene scores based on marker genes in the perturbed clusters using UCell^33^ and aggregated the scores by the gRNA type (Table S4). Next, we applied the dosage response curve (“drc”) package^34^ to model the relationship between Cluster 1 gene scores and *NANOG* expression, and between Cluster 2 gene scores and *OCT4* expression. We fitted both a linear model, describing gradual responses, and a Hill model, capturing switch-like behavior. Adopting the criteria used in a previous study^13^, we determined a response to be better fitted by the Hill equation if its Aikaike Information Criterion (AIC) was lower than that of the linear equation by more than 2 units (i.e., ΔAIC (Linear – Hill) > 2). Conversely, we classified it as better fitted by the linear equation if ΔAIC (Linear – Hill) ≤ 2 and the linear model’s p-value was < 0.05 (Figure 2A). We found that the cell state’s response to *NANOG* perturbations was more closely fitted by the linear model whereas the response to *OCT4* perturbations was more closely fitted by the Hill model (Figure 2B, Table S4). This suggests that decreasing *NANOG* dosage induces a gradual shift from pluripotency, while reducing *OCT4* dosage triggers a more abrupt and threshold-dependent exit from the ESC state. We then extended the analysis to fit individual DEG’s response to the perturbations. We found the majority of the DEGs exhibited Hill-like response to their respective perturbations, with a higher proportion observed in Cluster 2 DEGs’ response to *OCT4* (Figure 2C, Table S4), consistent with the overall cell state’s more switch-like response to *OCT4* than to *NANOG* (Figure 2B).

**Figure 2.**
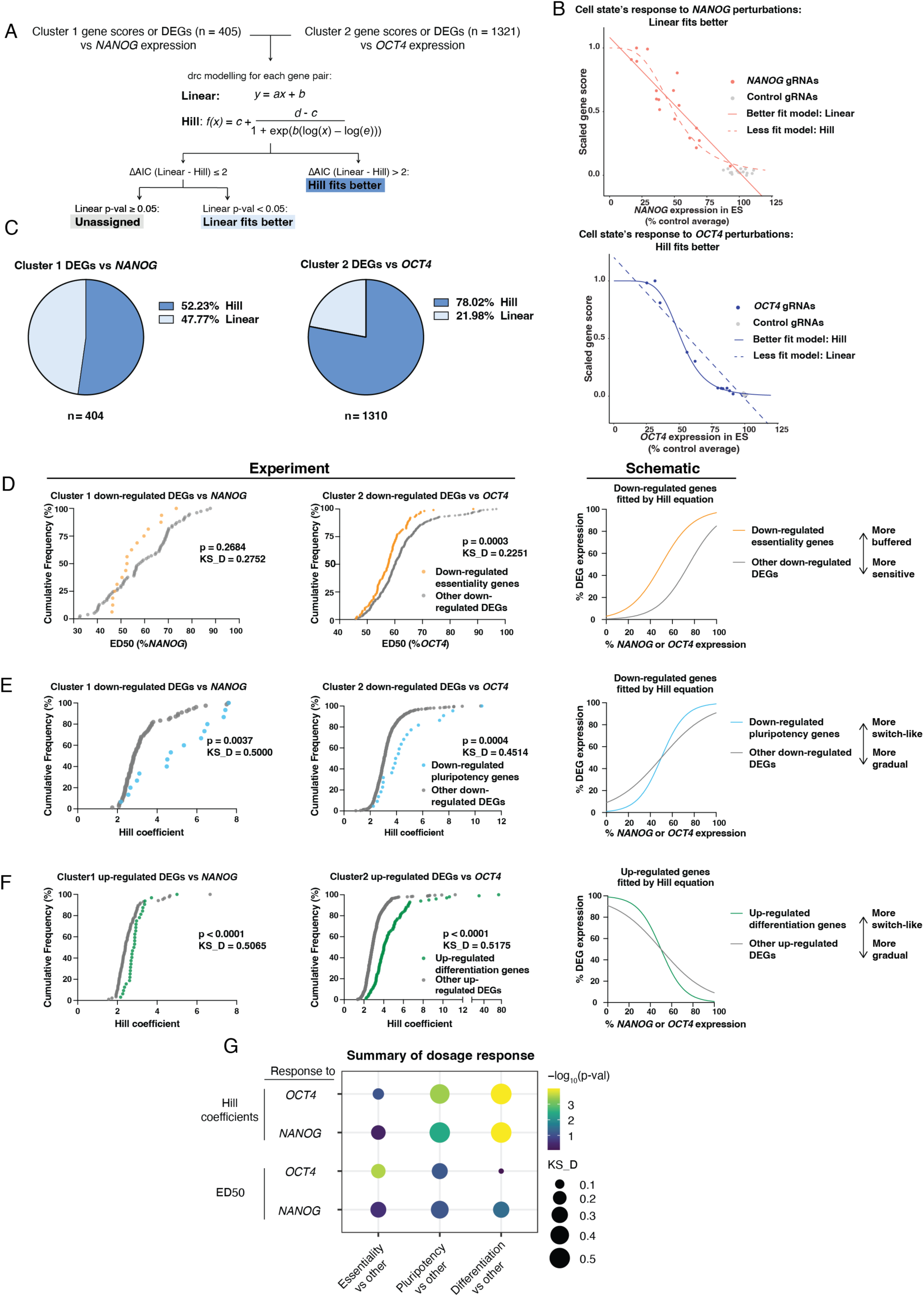
Dosage modeling of ESC Perturb-seq reveals distinct response patterns. A. Schematic of modeling workflow. B. Fitted models for the response of perturbation gene scores to *NANOG* or *OCT4* levels. Solid lines indicate the assigned model; dashed lines indicate the alternative model tested. C. Distribution of fitted models for Cluster 1 and Cluster 2 DEGs respectively. Genes without an assigned model were excluded (n = 1 for *NANOG* perturbations and 11 for *OCT4* perturbations). D. ED50 distribution showing essentiality genes were more buffered against *OCT4* perturbations than other down-regulated genes. E. Hill coefficient distribution showing pluripotency genes were more switch-like in response to *NANOG* and *OCT4* perturbations than other down-regulated genes. F. Hill coefficient distribution showing differentiation genes were more switch-like in response to *NANOG* and *OCT4* perturbations than other up-regulated genes. G. Summary of ED50 and Hill coefficient distribution for different gene sets in response to *NANOG* and *OCT4* perturbations. Statistical analysis for D-F: Two-sided Kolmogorov-Smirnov test.

Two parameters define the shape of the Hill curve: the Hill coefficient, which quantifies cooperativity and determines the steepness of the curve, and the effective dose 50 (ED50), which indicates the *NANOG* or *OCT4* dose at which each DEG reaches the midpoint of its differential expression. These parameters enable us to distinguish gradual vs switch-like response, and buffered vs sensitive response respectively. Since we observed enrichment of differentiation genes for up-regulation and enrichment of pluripotency and essentiality genes for down-regulation in the perturbed clusters (Figures 1D, S1G), we sought to understand their response dynamics by comparing their ED50s and Hill coefficients with other up- or down-regulated genes respectively. While DEGs related to differentiation and pluripotency did not significantly differ from other DEGs in their ED50 values, essentiality DEGs in Cluster 2 had significantly lower ED50 than other down-regulated DEGs, suggesting that essentiality genes were more buffered against *OCT4* perturbations (Figures 2D, G, S2B-C). On the other hand, whereas essentiality DEGs did not differ from other down-regulated DEGs in Hill coefficients, pluripotency and differentiation DEGs had significantly higher Hill coefficients than other down- or up-regulated DEGs, respectively (Figures 2E-G, S2A). This suggests that cell identity genes, those related to differentiation and pluripotency, respond to perturbations with accentuated switch-likeness, indicative of strong cooperative regulation.

### Perturbations of pluripotency TFs promote differentiation to both targeted and off-target lineages

The observation of the dosage-dependent effect of perturbing *NANOG* and *OCT4* on hESC self-renewal prompted us to investigate how the same perturbations influence differentiation. We induced definitive endoderm (DE) differentiation on hESCs infected with the same gRNA library. To capture the dynamics of the cell state transition, we harvested cells at the mid-point of ESC-to-DE differentiation (36 hours after induction) and referred to this dataset as DE Perturb-seq (Figure 3A). We observed a dosage-dependent decrease in cell representation with increasing perturbation levels, consistent with our observation in ESC Perturb-seq (Figures 1E, 3B). This trend was more pronounced in DE Perturb-seq, potentially reflecting additional selective pressure during differentiation. Seurat analysis showed that the majority of the cells followed a continuous differentiation trajectory: Cluster 0, the more differentiated group, expressed higher levels of DE markers *SOX17* and *GATA6*, and Cluster 1, the less differentiated group, expressed higher levels of ESC markers *NANOG* and *OCT4* (Figures 3C-D, Table S2). In addition, we observed a separate cluster, Cluster 2, which were enriched for *NANOG* or *OCT4* gRNAs (Figures 3C-D, S3A). It had low expression of both ESC and DE markers, reduced expression of essentiality genes (e.g., the metabolism enzyme *IDH1* and ribosomal component *RPS12*), and increased expression of genes associated with off-target lineages as represented by NE (e.g. *MAP2* and *GREB1L*; Figures 3E-F, S3B, Table S2). As perturbation levels increased from none (control gRNAs) to mild (Group 1 gRNAs) and then to moderate (Group 2 gRNAs), cells gradually shifted towards the more differentiated Cluster 0 (Figures 3G-H). As the perturbation levels increased from moderate (Group 2) to strong (Group 3), the fraction of cells in Cluster 0 decreased whereas the fraction of cells in Cluster 2 increased. Taken together, these data illustrate that different levels of *NANOG* or *OCT4* perturbations can result in divergent impact on DE differentiation, cell survival, and off-target differentiation.

**Figure 3.**
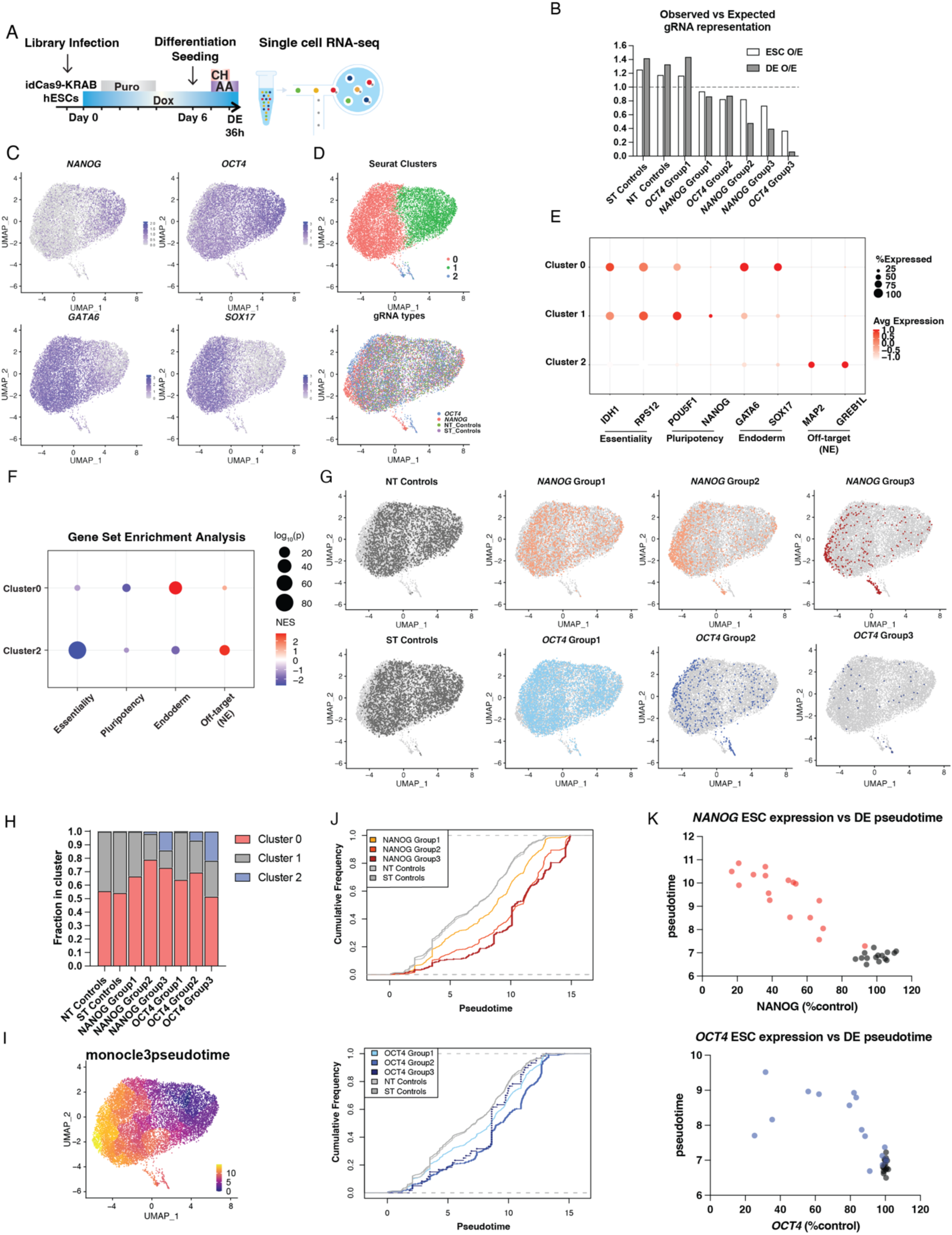
DE Perturb-seq shows diverging effects of perturbing *NANOG* and *OCT4* during differentiation. A. Schematic of the DE Perturb-seq. B. Representation for each perturbation group in ESC and DE Perturb-seq. C. UMAPs showing expression of ESC markers *NANOG* and *OCT4*, and DE markers *GATA6* and *SOX17*. D. UMAPs showing Seurat clustering and gRNA distribution. E. Cluster marker genes associated with essentiality, pluripotency, DE differentiation, and off-target (NE) differentiation. F. Summary of GSEA showing enrichment of genes associated with essentiality, pluripotency, DE differentiation, and off-target (NE) differentiation ranked by expression log2FC in Cluster 0 vs Cluster 1 and Cluster 2 vs Cluster 0 and 1. G. UMAPs showing distribution of gRNAs from different perturbation groups. H. Quantification of gRNA distribution in F. I. UMAP showing Monocle3 pseudotime. J. Cumulative frequency plot showing pseudotime for each perturbation group. K. Pseudotime trends aggregated by gRNA genotype.

We further examined the impact of perturbations on differentiation by quantifying the distribution of cells along the differentiation trajectory using Monocle3 pseudotime^35–37^ (Figure 3I). This analysis demonstrated that mild to moderate perturbations accelerated differentiation, whereas strong perturbations stopped improving or even reduced differentiation efficiency, particularly for *OCT4* (Figures 3J-K), indicating that the impact of decreasing pluripotency TF dosages on differentiation could be multi-directional instead of monotonic.

### Dosage of pluripotency TFs exhibits a non-monotonic effect on differentiation

The observation that different levels of pluripotency gene perturbations had divergent impact on differentiation prompted us to include non-monotonic models in our quantitative analyses. We chose the quadratic equation as a simple mathematical model to capture non-monotonic trends. To quantify differentiation and off-target effects, we defined a “differentiation” gene score using Cluster 0 marker genes, and an “off-target” gene score using Cluster 2 marker genes. We then modeled the relationship between these two scores and perturbation levels, measured by *NANOG* or *OCT4* expression at the ESC stage for each gRNA. To determine the best-fit model, we first compared the fit of linear and Hill equations. If the linear model provided a better fit than the Hill model, we next tested whether the quadratic model offered an even better fit as determined by ΔAIC (Linear – Quadratic) > 2 (Figures 4A), since the quadratic model is the linear model with the addition of a second order term. In contrast to the off-target gene score, which was best fitted by the Hill model in relation to *NANOG* and *OCT4* levels, the differentiation gene score was best fitted by the quadratic model (Figure 4B, Table S4). The differentiation gene score peaked at moderately reduced *OCT4* levels or strongly reduced *NANOG* levels, suggesting that partial reduction of pluripotency TFs may maximize differentiation efficiency.

**Figure 4.**
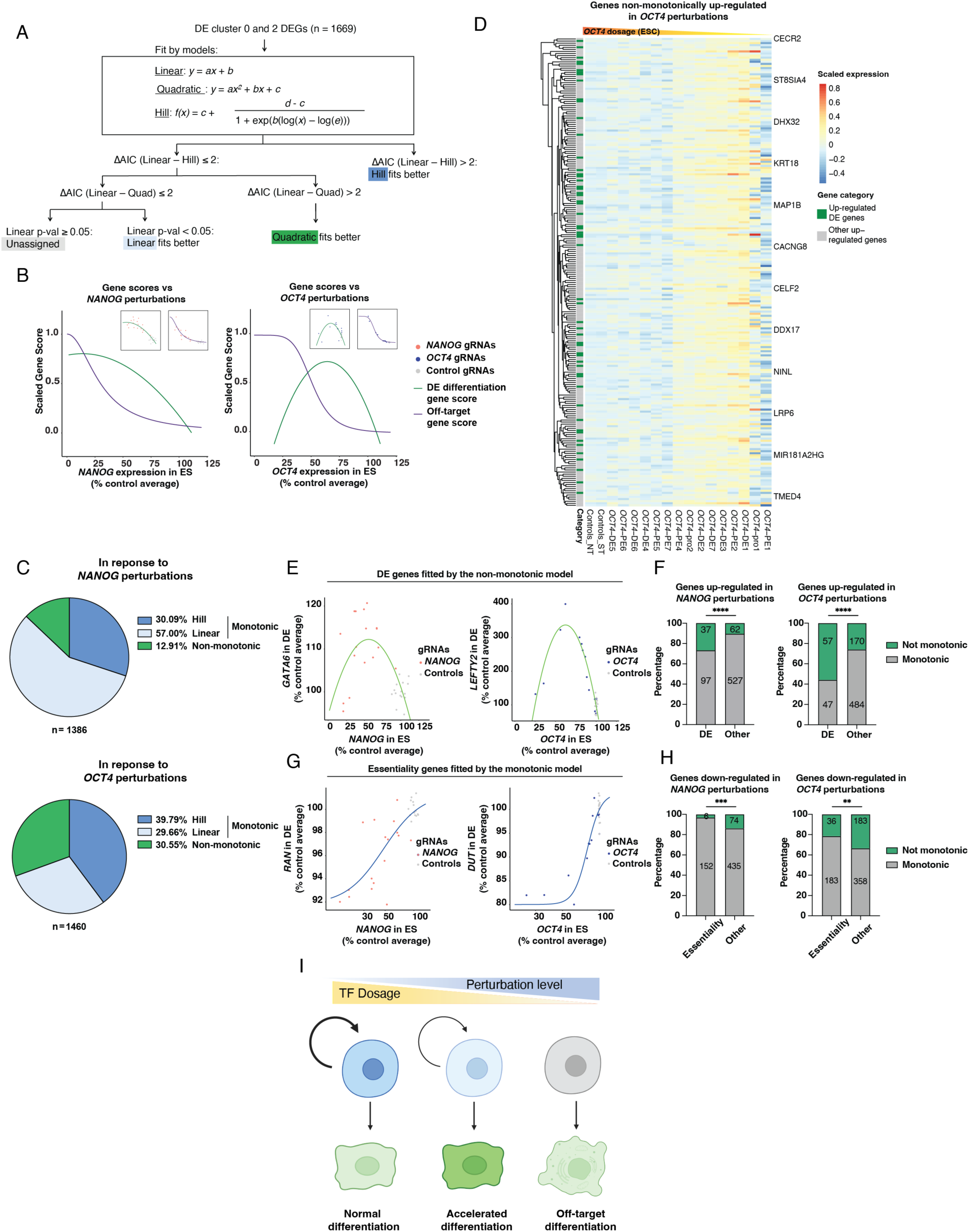
Dosage modeling of DE Perturb-seq identifies non-monotonic response of lineage specific genes. A. Schematic of modeling workflow. B. Fitted models for the response of differentiation (Cluster 0) and off-target (Cluster 2) gene scores to *NANOG* or *OCT4* levels; insets show values of individual gRNAs for each gene score before overlay. C. Distribution of fitted models for DE Cluster 0 and Cluster 2 DEGs’ response to *NANOG* and *OCT4* perturbations. Genes without an assigned model or assigned the non-monotonic model without change of direction within the experimental range were excluded (n = 283 for *NANOG* perturbations and 209 for *OCT4* perturbations). D. Heatmap with hierarchical clustering showing expression changes of DEGs non-monotonically up-regulated under *OCT4* perturbations. Color key indicates scaled expression. E. Examples of DE genes’ response curves fitted by the quadratic model. F. Distribution of monotonic and non-monotonic models for DE genes up-regulated under *NANOG* or *OCT4* perturbations. G. Examples of essentiality genes’ response curves fitted by the Hill model. H. Distribution of monotonic and non-monotonic models for essentiality genes down-regulated under *NANOG* or *OCT4* perturbations. I. Summary schematic. Statistical analysis for F and H: Fisher’s exact test, *p ≤ 0.05; **p ≤ 0.01; ***p ≤ 0.001; ****p ≤ 0.0001.

As an orthogonal validation of the modeling findings, we reanalyzed results from a previous CRISPRi screen^15^, conducted under DE differentiation in parallel with the ESC screen^23^ and utilizing the same gRNA library. The ESC and DE screen datasets allowed us to assess the effect of different levels of *NANOG* and *OCT4* perturbations (measured by depletion effect in the ESC screen) on differentiation (measured by gRNA distribution in cells expressing DE lineage marker in the DE screen). We found that gRNAs resulting in mild self-renewal impairment in the hESC screen were enriched in cells on the more differentiated side of the DE screen, whereas those with strong self-renewal effects were enriched on the less differentiated side of the DE screen (Figures S3C-E, Table S1). In contrast, gRNAs targeting validated DE enhancers^15^ did not exhibit the same dichotomy; most affected DE differentiation but had no impact on self-renewal (Figure S3E-F). These results reaffirmed the non-monotonic relationship between *NANOG/OCT4* dosage and differentiation efficiency.

In order to investigate genes contributing to different modes of cell state response, we further fitted the relationship between the expression of Cluster 0 and Cluster 2 DEGs and different levels of perturbations. We classified response curves best fit by the linear and Hill model as monotonic, and those best fit by the quadratic model as non-monotonic. We found that 87% of DEGs had monotonic relationships with *NANOG* perturbations and 13% had non-monotonic relationships. In contrast, 69% of DEGs had monotonic relationships with *OCT4* perturbations and 31% had non-monotonic relationships (Figure 4C, Table S4). This larger fraction of non-monotonic genes is consistent with the greater reduction of differentiation pseudotime (Figure 3K) and differentiation gene score (Figure 4B) at strong *OCT4* perturbations. These non-monotonic trends were also evident in the heatmap visualization of gene expression patterns (Figures 4D, S4A). To assess whether this non-monotonic response was specific to the DE Perturb-seq data, we included the quadratic equation in an updated modeling of the ESC Perturb-seq data. Only around 4-5% of DEGs showed non-monotonic responses to *NANOG* or *OCT4* perturbations in the ESC context (Figure S4B, Table S4), indicating that non-monotonic responses are a distinct feature of cells undergoing dynamic lineage transitions.

We evaluated the distribution of monotonic and non-monotonic responses in genes associated with cell identity and essentiality, as they were enriched for differential expression in DE Perturb-seq (Figures 3E-F). To facilitate the comparisons, we combined DEGs from Cluster 0 and Cluster 2 and categorized them as being up- or down-regulated in cells with *NANOG* or *OCT4* perturbations (Figure S5A). Consistent with the distribution of gRNAs and gene expression in these clusters, DE and NE genes were enriched among up-regulated DEGs, and essentiality genes as well as pluripotency genes were enriched among down-regulated DEGs (Figures S5B-E). Compared to other up-regulated DEGs, DE genes, representing on-target differentiation, were significantly enriched for non-monotonic relationships with both *NANOG* and *OCT4* perturbations, by approximately 3-fold (Figure 4E-F), whereas NE genes, representing off-target differentiation, were not (Figure S5F). Compared to other down-regulated DEGs, essentiality genes were significantly enriched for monotonic response to these perturbations, by 2-5 fold (Figures 4H-I), and pluripotency genes were not biased toward either type of response (Figure S5G). Taken together, these results demonstrate that non-monotonic responses to pluripotency factor perturbations are specific to differentiation into the targeted lineage.

## Discussion

In this work, we investigated how hESCs respond to varying dosages of *NANOG* and *OCT4* under self-renewal and differentiation conditions. The non-monotonic relationships we observed between many DE-specific genes and pluripotency TF dosages may help reconcile seemingly contradictory findings from previous studies. One study reported shRNA-mediated KD of *OCT4* reduced DE differentiation^22^. However, another study found that *OCT4* KD induced precocious expression of endodermal genes in hESCs, while *OCT4* overexpression decreased the expression of DE markers in DE differentiation^30^. In the case of *NANOG*, it has been reported that its KD impaired the specification of the DE lineage and its overexpression induced misexpression of early endoderm markers in hESCs^30^. However, *NANOG* overexpression was also shown to hinder DE differentiation^18^. Our findings suggest that these divergent outcomes may arise from the different degrees of *NANOG* and *OCT4* perturbations across experimental settings, which may either promote or inhibit differentiation depending on their position along a non-monotonic response curve. These findings emphasize the importance of understanding gene function through a continuum of expression levels, as binary perturbation approaches such as knockouts or single-dose knockdowns may overlook the non-linearity of downstream transcriptional and phenotypic consequences.

Gene dosage can be manipulated using different experimental systems. One approach is rapid protein depletion, such as with the auxin or dTAG systems^13,38^. Compared to dCas9-KRAB-mediated repression, protein depletion approaches offer greater precision in achieving the desired dosage and enable the examination of the immediate consequences of protein loss, making them particularly valuable when temporal resolution and fine dosage control are important to the research question. In comparison, the CRISPR-based system excels at scalability at the levels of both perturbations and readouts. At the perturbation level, it allows for the targeting of multiple genes without the need to engineer a separate tag for each gene. In addition, advances in predicting gRNA activity and designing attenuated gRNA allelic series further facilitate scalable titration of gene expression^39^. While we focused on CRISPRi, a dosage gradient can also been achieved by using CRISPR activation (CRISPRa) ^40,41^, as demonstrated by a recent study^42^. It would be particularly interesting to activate genes normally up-regulated during cell state transition at the pre-transition stage and study how the transcriptome responds to different dosages of activation. On the readout side, Perturb-seq simultaneously captures both the perturbation, represented by the gRNA, and its downstream responses across many cells in a single experiment. This makes it particularly well-suited for pooling cells receiving different perturbations for single-cell sequencing and detecting trends in a highly heterogenous cell population, such as differentiating cells. Notably, while we often assume that gene response direction remains constant across a dosage range, our observation of non-monotonic responses of lineage-specific genes to *OCT4* and *NANOG* perturbations during differentiation suggests that this assumption may need to be adjusted especially in dynamic systems. This underscores the importance of using methods capable of resolving cellular heterogeneity to fully capture gene dosage responses in cell state transitions.

Quantitative methods are essential to uncover patterns of gene dosage response in the experimental data. Here we applied the linear, Hill, and quadratic equations as a simple yet effective modeling approach to assess gene dosage response. This has allowed us to uncover trends consistent with biological intuition that have not been systematically characterized before. While our current analysis focused on individual gene responses, we envision building gene regulatory networks (GRNs) by modeling TF direct inputs to individual enhancers^43^ will further elucidate the mechanisms behind different dosage response patterns. For example, the predominant fit of cell identity genes to the Hill equation suggests these genes are regulated with high cooperativity; a network-based approach could elucidate the extent to which these genes receive input from multiple genes and regulatory elements. Similarly, non-monotonic genes may result from the indirect influence of opposing pathways, a hypothesis that can be tested by network analysis taking into account different feedback relationships. Together, these experimental and computational approaches can provide a framework for understanding gene dosage effects underlying cell state transitions not only in stem cell differentiation and early development, but also in broader health and disease transitions.

## Data availability

ESC and DE perturb-seq data have been deposited in the GEO database under the accession code GSE283614. RNA-seq of ESC and differentiation have been previously published under accession codes GSE213394 (DE36h differentiation) and GSE283612 (NE72h differentiation).

## Code availability

Custom scripts in this manuscript build on scripts published previously with modifications and can be accessed on GitHub: https://github.com/huangfulab/Code-for-ES-DE-modeling

## Author contributions

J.Y. and D.H. conceptualized the study, devised the experiments, and interpreted results. J.Y. performed most of the experiments and analyzed the results. H.S.C. performed dosage modelling. R.L. assisted with the single-cell experiments and analysis. M.A.B. and W.L. contributed to computational analysis and provided input for modeling work. J.Y. and D.H. wrote the manuscript; all authors provided editorial advice.

## Acknowledgements

We acknowledge the assistance from the following MSKCC Cores: Antibody & Bioresource, Flow Cytometry, and Integrated Genomics Operation (IGO). This study was funded in part by the National Institutes of Health grant U01HG012051 (D.H.), National Institutes of Health grant U01DK128852 (D.H.), Starr Tri-I Stem Cell Initiative #2023-006 (D.H.), NIH/NCI MSKCC Cancer Center Support Grant P30CA008748, Cycle for Survival (to IGO), and the Marie-Josée and Henry R. Kravis Center for Molecular Oncology (to IGO). Figure 4I was partially created with BioRender.com.

## Declaration of generative AI and AI-assisted technologies in the writing process

During the preparation of this work the authors used ChatGPT developed by OpenAI to improve the clarity of some sentences. The authors reviewed and edited the content as needed and take full responsibility for the content of the publication.

## Methods

### Culture of hESCs

Human ESCs were maintained in Essential 8 medium (E8; Thermo Fisher Scientific, A1517001) on tissue culture-treated polystyrene plates coated with 5 μg/mL vitronectin (Thermo Fisher Scientific, A14700) at 37 °C with 5% CO2. For regular maintenance, cells were passaged every 2-4 days, where they were treated with 0.5 mM EDTA (KD Medical, RGE-3130) in PBS for dissociation. For long term preservation, cells were dissociated, resuspended in E8 medium with 10% DMSO (Santa Cruz Biotechnology, sc-358801), and frozen in liquid nitrogen. For seeding cells, 10 μM Rho-associated protein kinase (ROCK) inhibitor Y-27632 (Selleck Chemicals, S1049) was added into the culture medium. Cell counting was performed on a Vi-CELL XR Cell Viability Analyzer (Beckman Coulter). Cells were routinely confirmed to be mycoplasma-free by the MSKCC Antibody and Bioresource Core Facility. This study used the idCas9–KRAB SOX17^eGFP/+^ HUES8 hESC line. The generation of this line was described in more details in Luo et al.^15^.

### Definitive endoderm differentiation

Definitive endoderm differentiation was performed as previously described^15^. Briefly, hESCs grown to ∼80% confluency in a “recovery passage” were washed with PBS and treated with TrypLE Select at room temperature for 3 min. After the treatment, TrypLE was removed, and hESCs were dissociated into single cells, resuspended in E8 medium, spun at 200g for 2 minutes, and resuspended in E8 medium with 10 μM ROCK inhibitor. 1 million cells were seeded per well on a 6-well plate (Fisher Scientific 0720080), or 0.3 million cells per well on a 12-well plate (Fisher Scientific, 0720082), coated with 5 μg/mL vitronectin. 24 hours after seeding, cells were washed with PBS and exposed to S1/2 medium containing 50 ng/mL Activin A (Bon-Opus Biosciences, C687-1MG) and 5 μM CHIR99021 (Tocris Bioscience, 4423). 24 hours later, the medium was changed to S1/2 containing only 50 ng/mL Activin A. After another 12 hours, cells were harvested for DE 36h time point. To make S1/S2 medium, MCDB131 medium (Thermo Fisher Scientific, 10372019) was supplemented with 1.5 g/L sodium bicarbonate (Research Products International, S22060), 1x Glutamax (Thermo Fisher Scientific, 35050061), 10 mM glucose (Sigma-Aldrich, G8769), and 0.5% BSA (LAMPIRE, 7500804).

### Perturb-seq gRNA library cloning

Cloning was performed individually for all 48 gRNAs following previously published methods^28,44,45^. For each gRNA, a pair of oligonucleotides (top: 5′-CACCG(N)_20_-3′; bottom: 5′-AAAC(N)_20_C-3′) were ordered from Eton Bioscience. 1 μL of each 100 μM oligonucleotides were mixed with 1 μL 10X T4 ligation buffer (NEB B0202S), 6.5 μL nuclease-free water (Thermo Scientific AM9938), and 0.5 μL T4 PNK (NEB M0201S) and incubated first at 37 °C for 30 minutes and at 95 °C for 5 minutes, followed by temperature ramp-down at 5 °C/min to 25 °C for annealing. The annealed oligonucleotides were diluted 1:200. 1 μL oligo duplex (diluted to 50 nM) were mixed with 50 ng BsmBI (Thermo Fisher Scientific, ER0451) digested CROPseq-Guide-puro plasmid (addgene: 86708), 1 μL T4 ligase (NEB M0202L), and 1x T4 ligase buffer (NEB B0202S) and incubated at room temperature for 20 minutes. The ligated plasmids were individually transformed into chemically competent Stbl3 *E.coli* (Invitrogen, C7373) followed by colony picking. Plasmids were extracted using the Zyppy Plasmid Purification Kit (Zymo Research, D4020), sequenced (Eton Bioscience), and mixed as into an equimolar pool to be packaged as lentivirus following transfection protocol described below.

### Lentivirus production and infection

500,000 293T cells were seeded in each well on a 12-well plate coated with 10 μg/mL collagen (Millipore Sigma, 08115) and cultured in DMEM media reconstituted with 15% Fetal Bovine Serum (Sigma Aldrich 12103C), 1x MEM Non-Essential Amino Acids (Thermo Fisher Scientific, 11140050), 1x Glutamax (Thermo Fisher Scientific, 35050061), and 1 mM sodium pyruvate (Sigma Aldrich, S8636). A day after seeding, cells were changed into fresh medium before transfection. For transfection in a 12-well, 0.1 μg vesicular stomatitis virus G (VSVG) envelope expressing plasmid pMD2.G (Addgene 12259), 0.4 μg lentiviral packaging vector psPAX2 (Addgene 12260), 1 μg lentiviral vector, and 3 μL JetPrime DNA transfection reagent were mixed with 75 μL JetPrime buffer (VWR, 89129924), incubated at room temperature for 10 minutes, and added drop wise to 293T cells. Medium was changed one day after transfection. Viral supernatant was collected 3 days after transfection, filtered through a 0.45 μM filter (Sigma Aldrich, SLHPR33RS), and stored at −80 °C. On the day of infection, virus supernatant was added to 1 mL E8 medium containing 10 μM ROCK inhibitor and 6 μg/mL protamine sulfate in a vitronectin-coated well on a 6-well plate before 75,000 hESCs resuspended in 1mL E8 medium containing 10 μM ROCK inhibitor were seeded on top. Antibiotic selection with 1 μg/mL puromycin (Sigma Aldrich, P8833) was performed 48 hours after seeding for 3 days. Experiment was scaled up or down as needed.

### ESC and DE Perturb-seq experiments

Approximately 225,000 cells and 450,000 cells in total, seeded in 6-well plates, were infected for the scRNA-seq experiment in ESC and DE respectively. For the experiment in ESC, one day after infection, 2 μg/mL doxycycline were added to the hESC culture until cell harvest on Day 6, with the addition of 1 μg/mL puromycin from Day 1 to Day 3 to select infected cells. Cells were passaged on Day 4 and harvested on Day 6. For the experiment in DE, 2 μg/mL doxycycline were added from Day 1 to cell harvest at DE 36h, with the addition of 1 μg/mL puromycin from Day 1 to Day 3 to select infected cells. Cells were seeded on Day 4 for “recovery passage” and seeded for differentiation on Day 6. For dissociation into single cells, adherent cells were washed with PBS, treated with TrypLE for 5 minutes at room temperature, gently triturated in FACS buffer, passed through 20 μM filters (Sysmex, 040042325), washed once with 0.5 mM EDTA and twice with PBS containing 0.04% BSA (LAMPIRE, 7500804). An aliquot of cell suspension was mixed 1:1 with Trypan Blue (Sigma Aldrich, T8154), loaded onto a cell counting slide (Invitrogen, C10228), and counted using a Countess Automated Cell Counter (Fisher Scientific). Then cells were loaded onto a 10x chromium controller v5.0 and processed with 10x Genomics Chromium Single Cell 3′ Reagent Kit v.3 following the manufacturer’s instructions (CG000204 Rev D). cDNA was assessed for quantity and quality control on a BioAnalyzer (Agilent) by the MSKCC Integrated Genomics Operation core. 10 μL cDNA was processed following the 10X protocol to generate the transcriptome library, and another 10 μL cDNA was used to generate the gRNA library. Briefly, cDNA was first amplified with primers overlapping the U6 promoter and R1 sequence on the captured gRNA transcript. Following clean-up with 1x SPRI Select (Beckman Coulter, B23317), the sample was then amplified with primer mix from the 10x Dual Index Kit TT Set A (PN-1000215) and cleaned up with 1x SPRI Select. The PCR condition and primer sequence are listed in Table S5. The library was checked for quantity and quality on a BioAnalyzer, and sequenced NovaSeq 6000 platform following the manufacturer’s guidelines by the MSKCC Integrated Genomics Operation core. The transcriptome libraries were sequenced for 400M and 600M reads for the ESC and DE experiment respectively. The gRNA libraries were each sequenced for 10M reads.

### Processing of Perturb-seq data

Alignment of scRNA-seq sequencing reads was performed using Chromium v3 analysis software Cell Ranger (version 3.1.0) pipeline “cellranger count”. “filtered_feature_bc_matrix” from the Cell Ranger output, containing both Gene Expression and CRISPR Guide Capture matrices, were processed by the Seurat package (v4.4.0). Quality control was performed to remove dead cells or multiplicates using the following cutoffs: 2500 < nFeature_RNA < 8000, nCount_RNA < 40000, percent.mt < 10 for the ESC experiment; 2000 < nFeature_RNA < 7000, nCount_RNA < 20000, percent.mt < 10 for the DE experiment. Cell cycle genes were regressed out. Furthermore, gRNA singlets (cells with exactly one type of gRNA) were selected using the following criteria: the gRNA with the highest UMI counts within a cell (“max gRNA”) had to have a UMI ≥ 5; the gRNA with the second highest UMI counts within that cell (“second max gRNA”) had to have a UMI < 5 and < 50% the UMI of the max gRNA. This analysis yielded 6076 and 12611 cells post-filtering for the experiment in ESC and DE respectively. PCA was calculated using top 2000 variable features. ESC and DE UMAPs were calculated using top 20 and 40 PCs respectively. Differentially expressed genes (DEGs) were performed using the FindMarker function using cutoffs |log_2_FC| > 0.25 and min.pct > 0.25. Up-regulated DEGs in each cluster are referred to as marker genes. For GSEA using ranked lists from scRNA-seq, the log_2_FC was calculated using the FindMarker function with only the min.pct cutoff. Pseudotime analysis was done by loading the DE Seurat object onto the Monocle3 package (v1.3.1) and choosing the top 2000 variable features as ordering genes. The root located on the most eastern (less differentiated) part of the UMAP was selected.

### Gene set selection for enrichment analysis

Based on RNA-seq data from DE and NE differentiation^15,23,31^, we defined DE and NE genes as those up-regulated in DE or NE respectively, and pluripotency genes as those down-regulated in both differentiation contexts. If a gene is up-regulated in both DE and NE, it is included in the DE gene set but not the NE gene set, to create a more inclusive set for assessing on-target differentiation and more stringent set for assessing off-target differentiation. In ESC Perturb-seq analysis, DE and NE genes were combined as differentiation genes. Essentiality genes were defined based on the integrated results of CRISPR-Cas9 essentiality screens performed across hundreds of cancer lines from DepMap version 22Q4^46^. The gene sets are listed in Table S3.

### CRISPR-seq of library-infected cells

Concurrent with the single-cell RNA-seq experiment, hESCs infected with the same gRNA library were seeded in a separate well, underwent the same antibiotics selection (puromycin for 3 days) but received no doxycycline treatment (“No Dox”). On Day 6, “No Dox” hESCs were pelleted, and their genomic DNA was extracted using the QIAGEN Blood & Cell Culture DNA Maxi Kit. The gRNAs were amplified using Herculase (Agilent Technologies, 600679) and primers targeting the U6 promoter and gRNAs scaffold on the CROP-seq backbone (forward primer: ACGATACAAGGCTGTTAGAGAGA; reverse primer: ACGGACTAGCCTTATTTTAACTTGC). The samples were sequenced for a maximum of 100,000 reads by CRISPRseq at the MSKCC Integrated Genomics Operation core. CRISPResso2 (http://crispresso.pinellolab.org/submission) was used for demultiplexing and quantification of barcode representation.

### Modeling of dosage-response curves

For input for modeling, UCell scores calculated using the selected DEGs (in the case of modeling gene scores) or normalized RNA counts from the Seurat Object (in the case of modeling individual gene expressions) were aggregated by taking the average value across cells containing the same gRNA. In ESC, the response of Cluster 1 gene score or DEGs to *NANOG* dosage was fitted using values from hESCs containing *NANOG* or control gRNAs, and the response of Cluster 2 gene score or DEGs to *OCT4* dosage was fitted using values from hESCs containing *OCT4* or control gRNAs. In DE, the response of Cluster 0 and Cluster 2 gene scores or DEGs to *NANOG* dosage was fitted using values from cells containing *NANOG* or control gRNAs, and the response to *OCT4* dosage was fitted in cells containing *OCT4* (excluding *OCT4*-PE3, one of the strongest *OCT4* gRNAs, as it was found in only 5 cells and was the only gRNA with a representation < 10) or control gRNAs. *NANOG* and *OCT4* dosage in both ESC and DE modeling was defined by the ESC data. The data was fitted to the linear model by the lm function in R and to a four-parameter Hill equation by the drm function in the drc package (version 3.0-1) utilizing the log-logistic function LL.4. The boundary conditions were set such that the upper limit of the Hill equation was fixed to the maximum aggregated expression value of the gene of interest (or gene score), and the lower limit was fixed to the minimum aggregated value, for more stabilized fitting of the Hill model.

In case of genes with non-monotonic perturbation effect, the data were fitted by the quadratic equation using the multivariable linear regression using lm. The quadratic formula was set up to the third degree, and with the parameter raw set TRUE to ensure interpretability. We confirmed that almost all DEGs assigned with the non-monotonic model exhibited non-monotonic trends within the experimental range (>97% for *NANOG* perturbations, and >99% for *OCT4* perturbations).

In ESC Perturb-seq, 1 Cluster 1 DEG (<1%) could not be assigned with a model in response to *NANOG* perturbations and 11 Cluster 2 DEG (<1%) could not be assigned with a model in response to *OCT4* perturbations. In DE Perturb-seq, 279 (17%) DEGs could not be assigned with a model under *NANOG* perturbations and 207 (12%) DEGs could not be assigned with a model under *OCT4* perturbations, possibly reflecting a higher level of gene expression heterogeneity in differentiation. These genes were excluded from pie charts showing model distribution and subsequent analyses.

---

**Figure S1.**
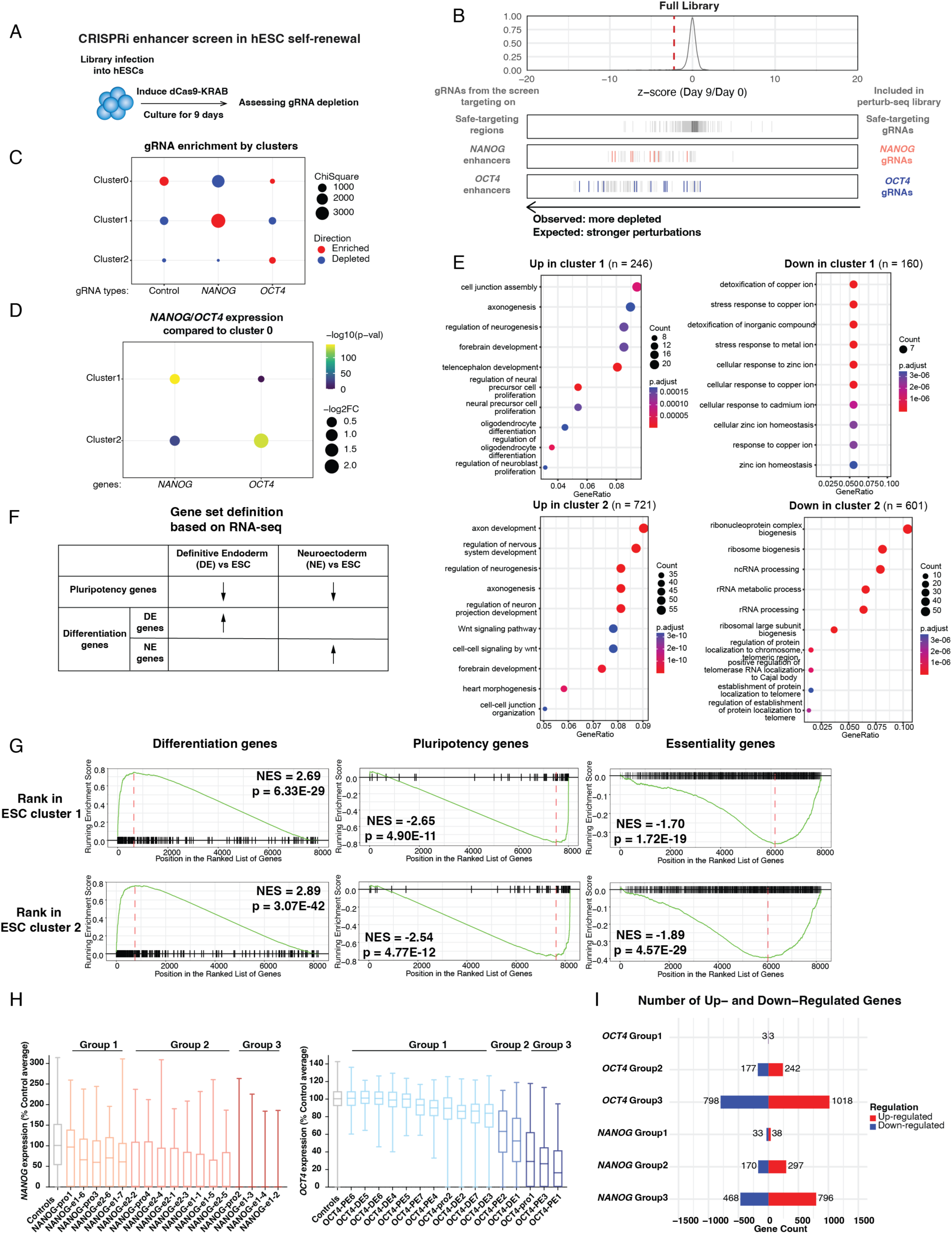
ESC Perturb-seq library design and results. A. Schematic of the previous CRISPRi enhancer screen in hESC self-renewal. B. Selection of gRNAs with different perturbation strengths based on screen results. C. gRNA enrichment in each cluster. Statistical analysis: Chi-squared test, p-value < 0.0001 for all comparisons. D. Differential expression of *NANOG* and *OCT4* in Cluster 1 and 2 relative to Cluster 0. Statistical significance (p < 0.05) is reached for *NANOG* in Cluster 1 and *OCT4* in Cluster 1 and 2. E. Gene ontology analysis of DEGs in Cluster 1 relative to Cluster 0 (upper panels) and DEGs in Cluster 2 relative to Cluster 0 (lower panels). Cutoffs for significant differential gene expression in E and F: |log2FC| > 0.25 and min.pct > 0.25. F. Schematic of GSEA gene set selection. Genes up-regulated during DE differentiation were excluded from the NE gene set to create a more stringent off-target gene set in DE Perturb-seq analysis. G. Individual GSEA plots for enrichment of differentiation, pluripotency, and essentiality gene sets ranked by expression log2FC in the Cluster 1 or Cluster 2 vs Cluster 0. H. Box plots showing the different distribution of *NANOG* or *OCT4* expression in cells receiving different levels of perturbations stratified into groups. I. Number of DEGs in gRNA groups with different targeting strengths. Cutoffs for differential gene expression: |log2FC| > 0.2 and min.pct > 0.25.

**Figure S2.**
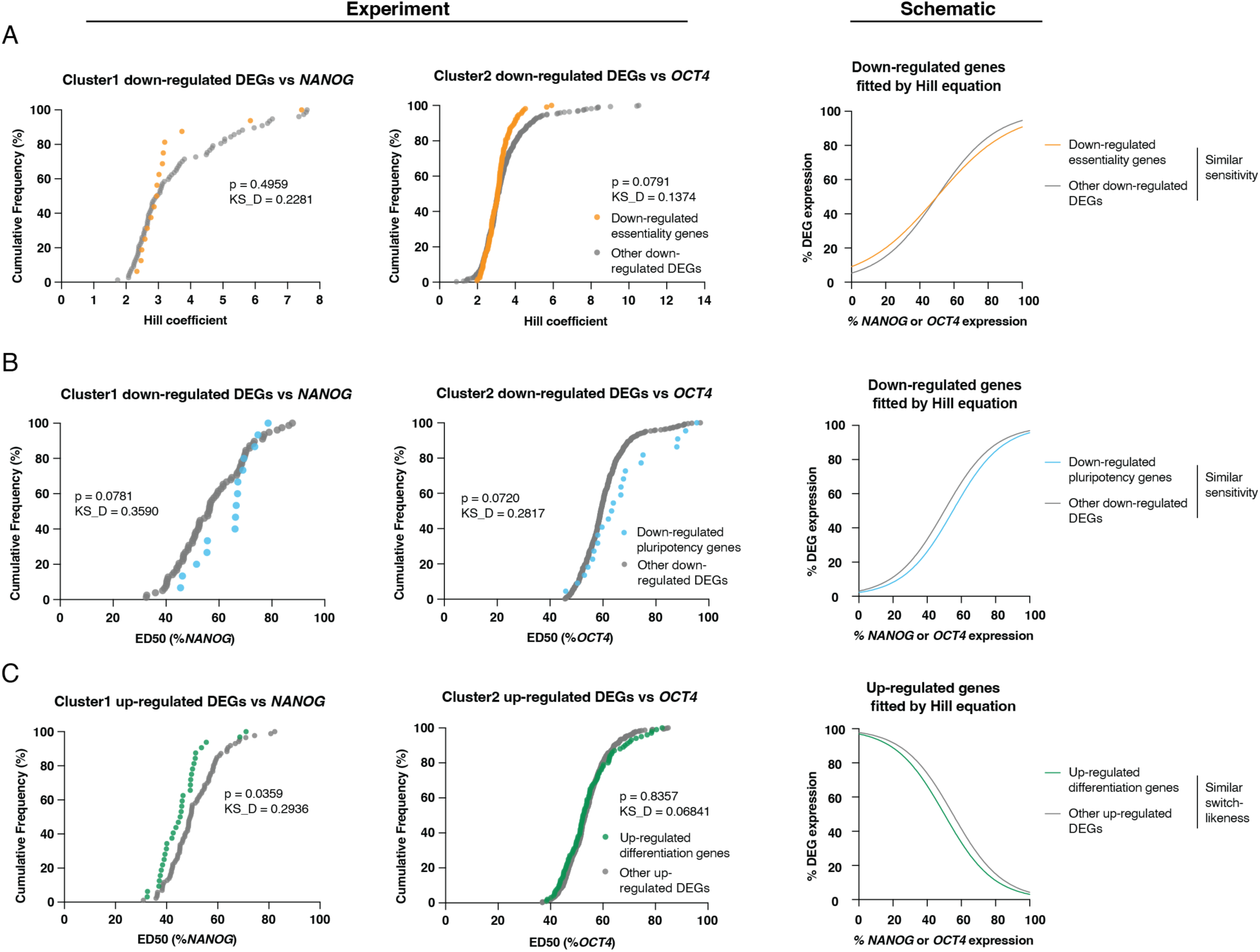
Indistinguishable response patterns from ESC dosage modeling. A. ED50 distribution showing essentiality genes were similarly switch-like as other down-regulated genes in response to *NANOG* and *OCT4* perturbations. B. Hill coefficient distribution showing pluripotency were similarly sensitive as other down-regulated genes in response to *NANOG* and *OCT4* perturbations. C. Hill coefficient distribution showing differentiation genes were similarly sensitive as other up-regulated genes in response to *NANOG* and *OCT4* perturbations. Statistical analysis for A-C: Two-sided Kolmogorov-Smirnov test.

**Figure S3.**
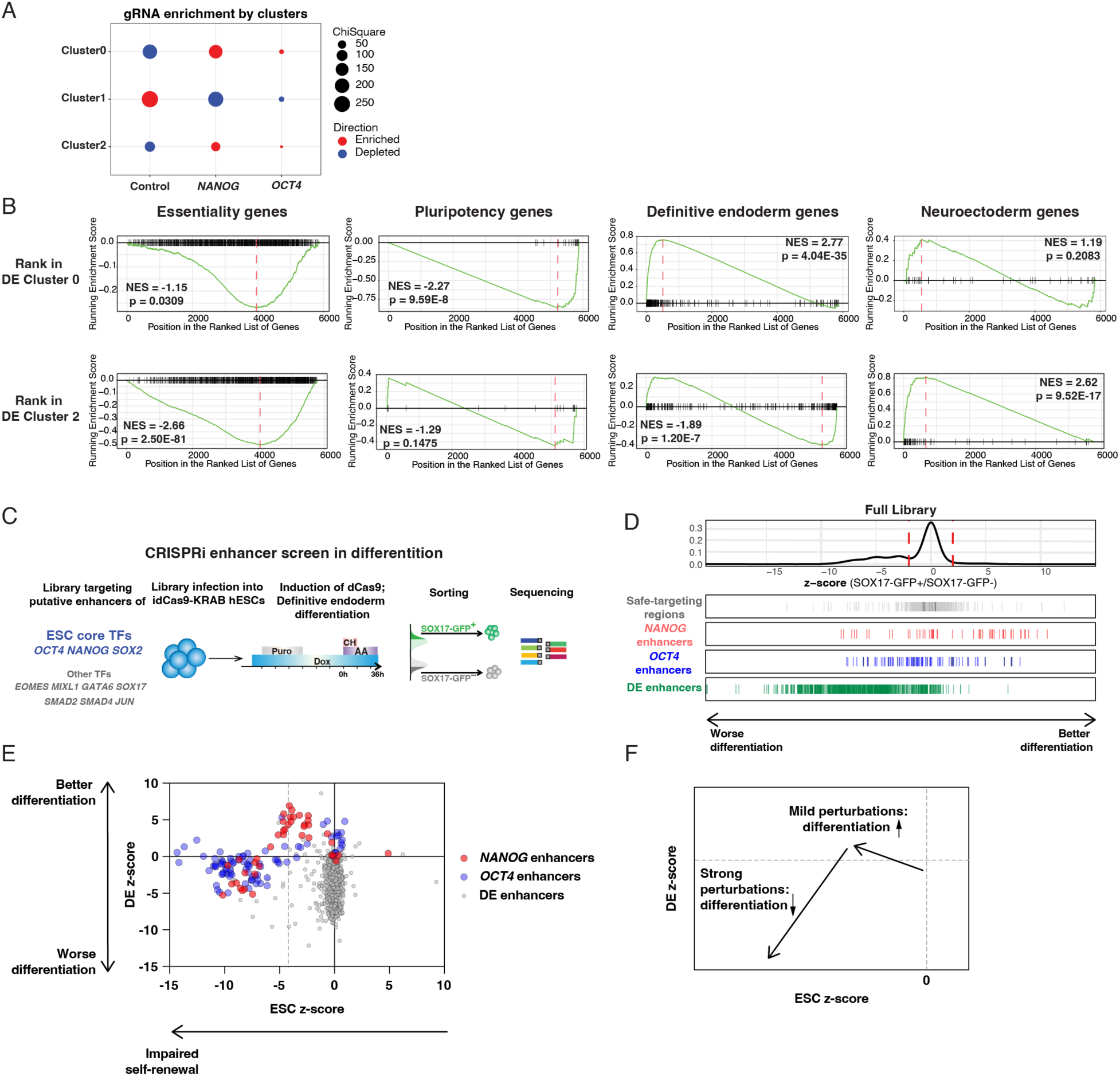
DE Perturb-seq differential gene expression and comparison with CRISPRi screen. A. gRNA enrichment in each cluster. Statistical analysis: Chi-squared test, p-value < 0.0001 for all comparisons, except for *OCT4* gRNA enrichment in Cluster 0 (p = 0.0004) and Cluster 2 (p = 0.0054). B. Individual GSEA plots for enrichment of essentiality, pluripotency, DE differentiation, and off-target (NE) differentiation gene sets ranked by expression log2FC in the DE Cluster 0 (“more differentiated”) vs Cluster 1 (“less differentiated”), and Cluster 2 (“off-target”) vs Cluster 0 and 1. C. Schematic of our previous CRISPRi enhancer screen in hESC definitive endoderm differentiation. D. The effects of individual gRNAs on differentiation as shown by the z-scores of log_2_FC(SOX17-GFP^+^ normalized reads/ SOX17-GFP^-^ normalized reads), normalized to ST gRNAs. E. Scatter plot showing the effect of individual gRNAs targeting ESC and DE enhancers in the ESC screen and the DE differentiation screen. Each symbol represents an individual gRNA. F. Schematic summarizing the trends observed in E.

**Figure S4.**
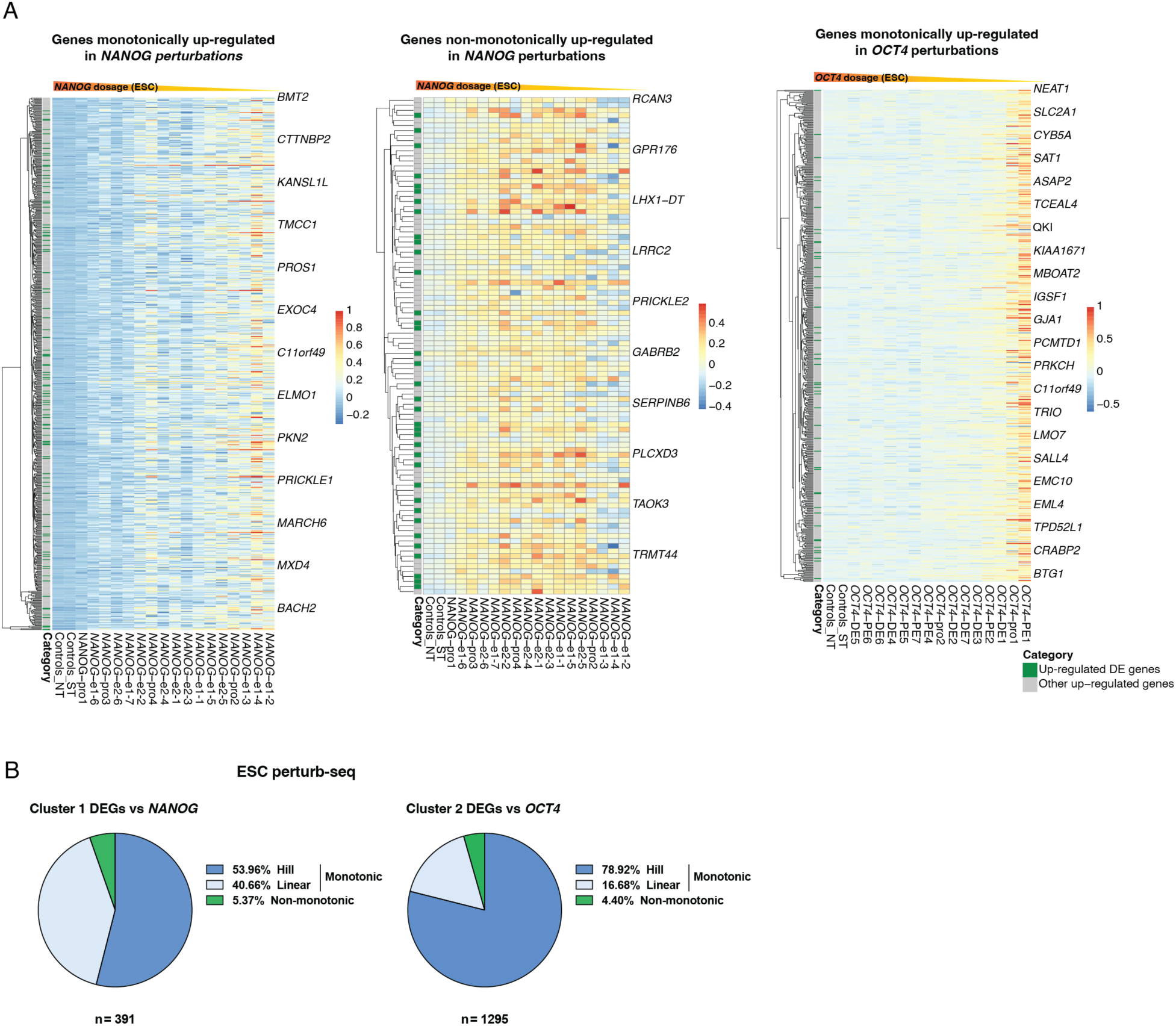
Monotonic and non-monotonic responses in ESC and DE Perturb-seq. A. Heatmap with hierarchical clustering showing expression changes of DEGs monotonically up-regulated under *NANOG* perturbations, non-monotonically up-regulated under *NANOG* perturbations, and monotonically up-regulated under *OCT4* perturbations. Color key indicates scaled expression. B. Distribution of fitted models, including the quadratic model, for ESC Cluster 1 and Cluster 2 DEGs’ response to *NANOG* and *OCT4* perturbations respectively. Genes without an assigned model or assigned the non-monotonic model without change of direction within the experimental range were excluded (n = 14 for *NANOG* perturbations and 26 for *OCT4* perturbations).

**Figure S5.**
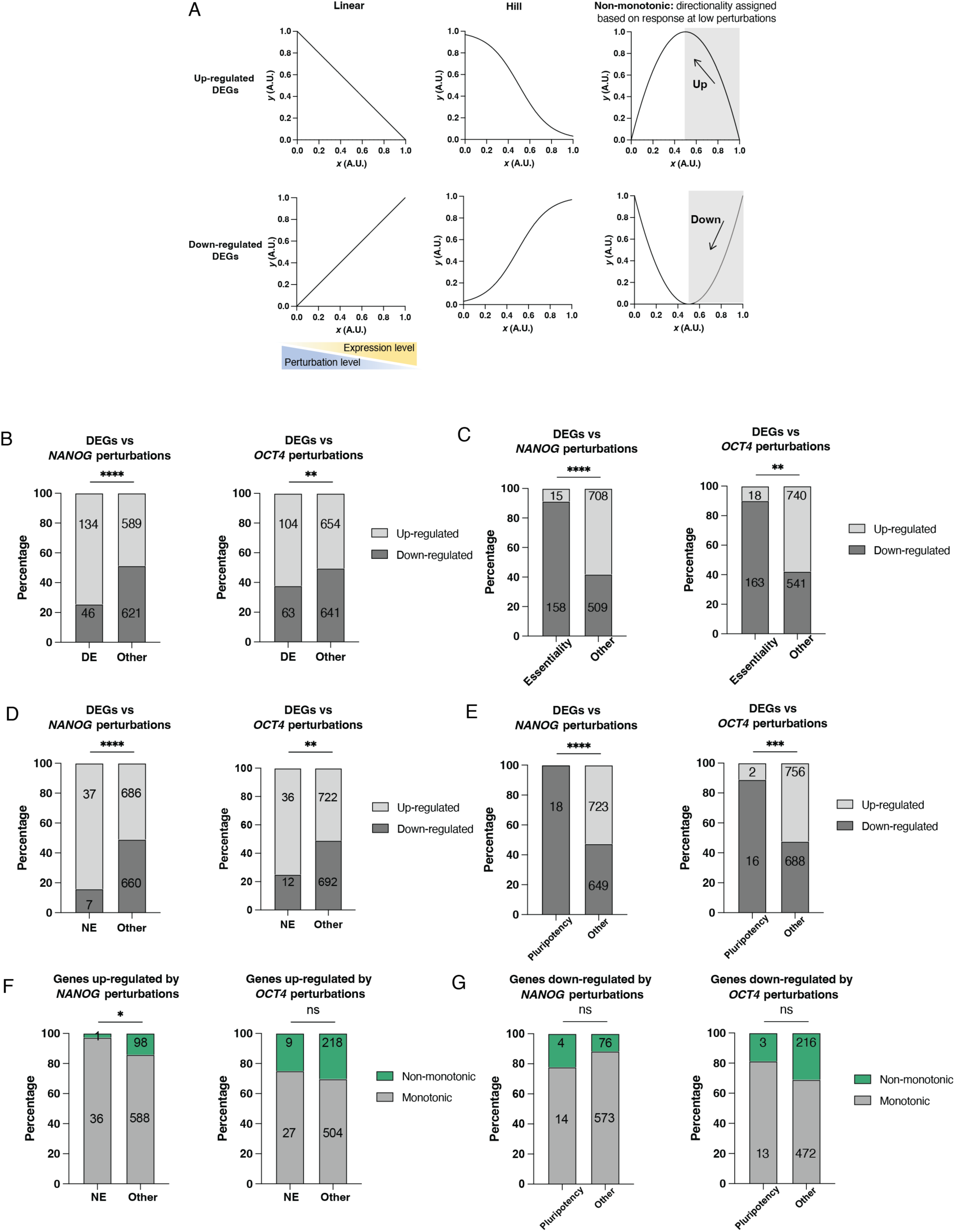
Response patterns for different gene sets in DE Perturb-seq. A. Schematics showing categorization of genes as being up-regulated or down-regulated by *NANOG* and *OCT4* perturbations. For genes with non-monotonic responses, we assigned their directionality based on the response at low perturbation levels, before any change in direction occurred (indicated by grey shading). B. Distribution of DE genes for being up-regulated or down-regulated under *NANOG* or *OCT4* perturbations. C. Distribution of essentiality genes for being up-regulated or down-regulated under *NANOG* or *OCT4* perturbations. D. Distribution of NE genes for being up-regulated or down-regulated under *NANOG* or *OCT4* perturbations. E. Distribution of pluripotency genes for being up-regulated or down-regulated under *NANOG* or *OCT4* perturbations. F. Distribution of monotonic and non-monotonic models for NE genes up-regulated under *NANOG* or *OCT4* perturbations. G. Distribution of monotonic and non-monotonic models for pluripotency genes down-regulated under *NANOG* or *OCT4* perturbations Statistics: Fisher’s exact test, *p ≤ 0.05; **p ≤ 0.01; ***p ≤ 0.001; ****p ≤ 0.0001.

